# A critical test of deep convolutional neural networks’ ability to capture recurrent processing in the brain using visual masking

**DOI:** 10.1101/2022.01.30.478404

**Authors:** Jessica Loke, Noor Seijdel, Lukas Snoek, Matthew van der Meer, Ron van de Klundert, Eva Quispel, Natalie Cappaert, H. Steven Scholte

## Abstract

Recurrent processing is a crucial feature in human visual processing supporting perceptual grouping, figure-ground segmentation, and recognition under challenging conditions. There is a clear need to incorporate recurrent processing in deep convolutional neural networks (DCNNs) but the computations underlying recurrent processing remain unclear. In this paper, we tested a form of recurrence in deep residual networks (ResNets) to capture recurrent processing signals in the human brain. Though ResNets are feedforward networks, they approximate an excitatory additive form of recurrence. Essentially, this form of recurrence consists of repeating excitatory activations in response to a static stimulus. Here, we used ResNets of varying depths (reflecting varying levels of recurrent processing) to explain electroencephalography (EEG) activity within a visual masking paradigm. Sixty-two humans and fifty artificial agents (10 ResNet models of depths - 4, 6, 10, 18 and 34) completed an object categorization task. We show that deeper networks (ResNet-10, 18 and 34) explained more variance in brain activity compared to shallower networks (ResNet-4 and 6). Furthermore, all ResNets captured differences in brain activity between unmasked and masked trials, with differences starting at ∼98ms (from stimulus onset). These early differences indicated that EEG activity reflected ‘pure’ feedforward signals only briefly (up to ∼98ms). After ∼98ms, deeper networks showed a significant increase in explained variance which peaks at ∼200ms, but only within unmasked trials, not masked trials. In summary, we provided clear evidence that excitatory additive recurrent processing in ResNets captures some of the recurrent processing in humans.

**Significance statement:** The challenge of modeling recurrent processes is not trivial and the operationalization of recurrent processing is highly contested. In this paper, we tested the ability of deep residual networks (ResNets) to explain recurrent processes in the human brain. Though ResNets are feedforward networks, they have been shown to equate operations in recurrent neural networks. In this study, we show that deeper networks explained more variance in brain activity than shallower networks. However, all networks still performed far from the noise ceiling. Thus, we conclude that recurrent processing in ResNets captures a form of recurrent processing in humans though other types of recurrent processing (inhibition, multiplicative) that are not present in current regular deep neural networks (alexnet, cornet, resnet) are necessary for building better visual models.

## Introduction

Deep convolutional neural networks (DCNNs) are currently the best mechanistic models of object recognition and best at predicting human neural visual processing dynamics (Kietzmann, McClure, et al., 2019; Kriegeskorte, 2015; Yamins & DiCarlo, 2016). However, DCNNs have also been critiqued for lacking crucial biological features, such as recurrent processing ((Kietzmann, Spoerer, et al., 2019; van Bergen & Kriegeskorte, 2020). Feedforward and recurrent processing are two modes of visual processing in biological brains (V. A. Lamme & Roelfsema, 2000). Though there is no strict separation between both modes, the distinction of both modes has been made spatio-temporally from electrophysiology, lesion studies, and neuropharmacological interventions (V. A. Lamme et al., 1998). Feedforward processing is a rapid set of computations evoked by sensory information, also known as bottom-up processing (Serre et al., 2007; Thorpe et al., 1996). Recurrent processing sets in right after the initial feedforward computations, and is known to include both lateral and top-down processing. In the context of object recognition, recurrent processing is believed to be essential for recognizing noisy, occluded objects (Rajaei et al., 2019; Spoerer et al., 2017; Tang & Kreiman, 2017), perceptual-grouping (Roelfsema, 2006) and figure-ground segmentation (Fahrenfort et al., 2007; Scholte et al., 2008). Its importance in human visual processing has led researchers to assert the lack of recurrent processes in deep convolutional neural networks (DCNNs) as a crucial limitation (Kietzmann, Spoerer, et al., 2019; Kreiman & Serre, 2020).

Consequently, researchers have attempted to incorporate recurrent processes in the architecture of DCNNs. This evoked discussions on different ways to model recurrent processes as researchers from separate disciplines have modeled recurrent processing differently. Physiologists who are primarily concerned with biological realism have modeled recurrent processing as an interaction between feedforward and feedback signals, influenced by neurons’ feature preferences and asymmetry of feedforward and feedback signals (Mély et al., 2018; Roelfsema et al., 2002). Others, mainly from the field of computer vision, modeled recurrent processing as a summation between feedforward and feedback signals (Kietzmann, Spoerer, et al., 2019; Kubilius et al., 2019; Tang et al., 2018). Regardless of approach, these different attempts at incorporating recurrent signals in models made clear that recurrent processing improves model performance - especially when the task at hand is challenging. Thus, there is a consensus on a necessity to incorporate recurrent processing in DCNNs; however, the appropriate level of complexity in approximating recurrent signals during the task of object recognition remains unresolved. One of the simplest models for recurrent processing (as proposed by the graduate student of the founder of computational neuroscience, Tomaso Poggio, and himself) is a *deep residual network* (referred to as “ResNets” here; (He et al., 2016b). The proposition of using ResNets as a model for recurrent processing follows from the observation that improvement in DCNNs performance over the years came mainly from incorporating additional layers in the network architecture, and these additional layers mimicked recurrent processes in primate brains (Liao & Poggio, 2016). Furthermore, Liao and Poggio have shown that ResNets’ computations are equivalent to unrolled time steps of recurrent computations in recurrent neural networks (RNNs); leading the authors to the conclusion that “moderately deep RNNs are a biologically-plausible model of the ventral stream in visual cortex”. Therefore, in our study, we used a family of ResNet models of different depths (ResNet-4, 6, 10, 18, 34) as proxies for varying levels of recurrent processing, to model recurrent signals in the human visual system. For each ResNet, we initialized and trained 10 seeds of the network, resulting in 50 artificial agents that we treat akin to animal models (Scholte, 2018).

Earlier studies have shown that task difficulty is an important factor for the occurrence of recurrent processes (Groen et al., 2018; Kar et al., 2019; Rajaei et al., 2019; Spoerer et al., 2017; Tang et al., 2018). In an earlier study, Seijdel, Loke et al. (2020) has similarly shown that image complexity predicts the occurrence of recurrent signals. Therefore, in the context of an object recognition task, image complexity could also be a factor of task difficulty - the higher the image complexity, the more difficult the task becomes. The current study complements the results from Seijdel, Loke et al. (2020) by testing the ability of deep residual networks to explain the different amounts of recurrent processing in neural data evoked through varying image complexity. We embedded the same target object in backgrounds with varying amounts of complexity and used visual masking to disrupt the occurrence of recurrent processes (Fahrenfort et al., 2007; Lamme et al., 2002). We posit that the combination of target objects from natural images on artificial backgrounds gives this experiment a good balance between naturalistic image qualities and experimental control. The expectation was that all networks, regardless of depth, would explain the same amount of variance in the brain for masked trials (when we disrupt recurrent processing); whereas deeper networks would be able to explain more variance of brain activity for unmasked trials (when visual processing is left unaffected), due to the approximation of recurrent processes in deeper layers. We found that deeper networks (ResNets-10, 18 and 34) indeed explained more variance of brain activity than shallower networks (ResNets-4 and 6). Furthermore, all ResNets explained more brain activity in unmasked than masked trials with differences between masking conditions starting as early as ∼98ms. In favor of replicability of model performance and variance estimation in ResNets (Mehrer et al., 2020), we trained 10 different seeds of each ResNet model for fitting neural data. These 50 ResNet models are made available at https://osf.io/hcj27/.

## Materials and methods

The subjects, stimuli and experimental paradigm are identical to the ones in Seijdel, Loke et al. (2020). However, for the ease of reading, we will also briefly describe them here. The human behavioral data has also been presented in Seijdel, Loke et al. (2020). In our results section, we summarized the human behavioral data and presented it against new networks behavioral data.

### Human Subjects

A total of 62 participants (45 females, 18-35 years old) took part in our experiment. Data from one participant was excluded due to the wrong placement of electrodes I1 and I2, and two other participants were excluded due to technical errors causing missing trials (because only one trial was obtained per stimulus, missing trials can not be used for a representational similarity analysis, see *Analysis: Representational Similarity Analysis (RSA)*.

### Stimuli

For the categorization task, we used a total of 120 unique images from five categories (i.e., 24 unique objects per category). These categories are bird, cat, fire hydrant, frisbee, and suitcase. The images were curated from several online databases - SUN (Xiao et al., 2010),

Microsoft COCO (Lin et al., 2014), Caltech-256 (Griffin et al., 2007), Open Images V4 (Kuznetsova et al., 2018), and LabelMe (Russell et al., 2008). The selected images underwent several preprocessing steps. First, we cropped the images into 512×512 pixels and converted them to grayscale. Second, we manually extracted the target objects in the images and then re-pasted the target objects onto one of four backgrounds. The first type of background is a uniform gray color, referred to as the “segmented” condition. The second type of backgrounds was generated by phase scrambling the background of the original images (after target object removal). The backgrounds differed in complexity as determined by two values - Spatial Coherence (SC) and Contrast Energy (CE) (Scholte et al., 2009). Forty images with the lowest SC and CE values were labeled as “low complexity” images; forty images with the highest SE and CE values were labeled as “high complexity”; and the remaining forty images in the middle range were labeled as “middle complexity” images. A previous study in our lab (Groen et al., 2018) has shown the validity of SC and CE values as an index of image complexity. Altogether, with 120 objects embedded in four different backgrounds, we have 480 unique stimuli.

### Experimental design

Participants performed a five-choice categorization task. The trial sequence is illustrated in Figure 1. The complete task consisted of 960 randomized trials (480 unique stimuli presented unmasked and masked), equally divided between visual masking, object category, and background complexity conditions. The trials were grouped into eight blocks of 120 trials with a one-minute break between each block (though participants were allowed a longer break if necessary).

**Figure 1.**
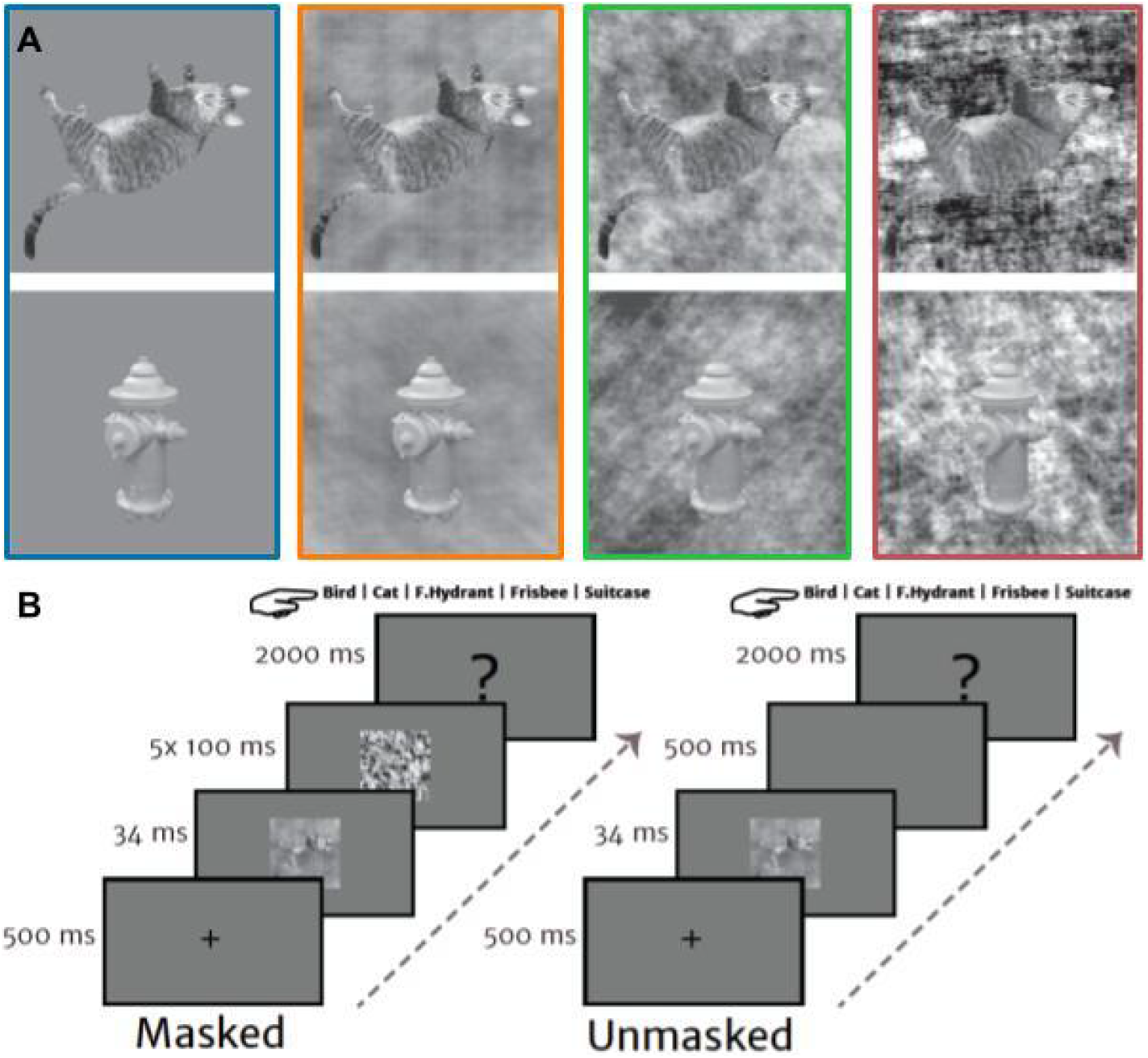
Stimuli and experimental paradigm. A) Exemplars of two categories (cat, fire hydrant) from each complexity condition. Backgrounds were either uniform (segmented; blue), or had low (orange), medium (green) or high (red) complexity values. B) Experimental design. On masked trials, the stimulus was followed by a dynamic mask (5×100 ms); on unmasked trials, the stimulus was followed by a blank screen (500 ms). Subsequently, participants were asked to categorize the target object. Figure taken from Seijdel, Loke et al. (2020).

The task was programmed in Presentation (Version 18.0, Neurobehavioral Systems, Inc., Berkeley, CA, www.neurobs.com) and presented on a 23-inch ASUS TFT-LCD display with a spatial resolution of 1920*1080 pixels and refresh rate of 60 Hz. Participants were seated approximately 70cm from the screen. The lights in the room were dimmed and kept constant between participants.

### Deep Residual Neural Networks

A family of ResNets was selected: ResNet-4, 6, 10, 18, 34. The numbers in the model names indicate the models’ total number of convolution and pooling layers. For each model, 10 different initializations were used as each initialization provides a certain amount of variance in its internal representations (Mehrer et al., 2020). Each of these initialized networks was trained with the ImageNet Large Scale Visual Recognition Challenge 2012 (ILSVRC) dataset and fine-tuned to the five object categories with a separate dataset from Microsoft COCO. For the initial training, we used a learning rate of 0.1, with a learning rate decay of 0.1 every 30 epochs. We used a stochastic gradient descent optimizer with a momentum of 0.9. All networks were trained for 150 epochs. For fine-tuning, we replaced the final fully-connected layer and retrained the weights for all layers. We used 13,648 training images and 584 test images from five categories for fine-tuning the network. In regards to fine-tuning hyperparameters, we used a learning rate of 0.001, with a learning rate decay of 0.1 every 7 epochs. We also used a stochastic gradient descent optimizer with a momentum of 0.9. By 40 epochs, the fine-tuning validation performance reached a plateau. All ResNets training, fine-tuning and feature extraction was performed in PyTorch (Paszke et al., 2019).

### EEG data acquisition and preprocessing

EEG recordings were made with a Biosemi 64-channel Active Two EEG system (Biosemi Instrumentation, Amsterdam, The Netherlands, www.biosemi.com) at a sample rate of 1024 Hz. A standard 10-10 electrode placement was used. As we were more interested in visual processing, electrodes F5 and F6 were moved to the occipital region and used as electrodes I1 and I2. Four external electrodes were used to record eye-movement artifacts. Preprocessing was performed in MNE-Python (Gramfort et al., 2013) using the following steps: (1) raw data was re-referenced to the average of left and right electrodes on the mastoids, (2) high-pass (0.1Hz) and low-pass (30 Hz) filters were applied, (3) Independent Component Analysis (ICA; Vigario et al., 2000) was performed to identify and remove remainder artifact components, specifically eye-movements and eye-blinks, (4) data were segmented into epochs from -100 to 600ms relative to stimulus onset, (5) baseline correction was applied to the 100ms prior to stimulus onset, (6) data were transformed to current source density responses (Perrin et al., 1989), (7) multivariate noise normalization was applied (Guggenmos et al., 2018).

### Analysis: Representational Similarity Analysis (RSA)

We examined both brain activity and internal representations of the ResNets using Representational Similarity Analysis (RSA; Kriegeskorte et al., 2008). RSA allowed us to test our hypotheses through the framework of representational geometry. We stored the pairwise distances between the activations of our experimental stimuli in representational dissimilarity matrices (RDMs). Importantly, these RDMs abstract away from the original measurement units and allow for comparisons across different domains - in our paper, between human brain activity and DCNNs activity.

From the EEG measurements, we obtained a RDM for every time sample from -100ms to 600ms relative to stimulus onset; this amounts to 180 RDMs per trial. The RDMs were computed based on activity from 22 posterior electrodes (Iz, I1, I2, Oz, O1, O2, POz, PO3, PO4, PO7, PO8, Pz, P1, P2, P3, P4, P5, P6, P7, P8, P9, P10). The electrodes were selected based on a previous study on the relationship between recurrent processing and image complexity (Groen et al., 2018).

From the DCNNs, we obtained RDMs from all convolutional layers. Similar to the analysis in Storrs et al. (2020), we transformed the activations using 100 principal components instead of using raw activations. These components were obtained with a separate dataset (*n*=2986). The transformed activations were then used to compute the pairwise (dis)similarity. The pairwise dissimilarity was computed as (1 − *Pearson correlation*) of the pattern responses. In the following analyses involving the RDMs, only the upper triangle is used (excluding the diagonal).

We used ResNets RDMs as predictors to explain EEG RDMs in a regression model. This regression was performed with non-negative least squares (NNLS) (Khaligh-Razavi & Kriegeskorte, 2014; Kaniuth & Hebart, 2021). The RDMs used in our regression models were made by trials averaged within the same object categories to increase the signal-to-noise ratio and place a larger focus on object categorization. After averaging trials within object categories, we obtained 20×20-sized RDMs - 5 object categories x 4 background conditions. We also performed trial averaging on the ResNets’ activations within object categories. The regression for each ResNet model is dependent on the number of layers in the model (see Figure 2). For example, with ResNet-6, we would build five different regression models. For the first model, we regressed only the first convolutional layer onto the EEG RDMs; for the second model, we regressed the first and second convolutional layers; this process continued until we included all five convolutional layers in ResNet-6. We performed 50 cross-validations for each model. The cross-validations folds were performed similarly as in Storrs et al. (2020). In every cross-validation, we reserved 15 (randomly chosen) test participants and 12 (randomly chosen) test stimuli per category (12 × 5 categories x 4 conditions = 240 trials in total), while fitting the NNLS model on 44 participants and 12 train stimuli. For each model, we computed a R^2^ value based on 15 test participants and 12 test stimuli. We also computed the upper and lower bounds of the noise ceiling by taking the averaged correlation of each test participant’s RDM with the RDM averaged across all test participants (upper bound), and taking the averaged correlation of each test participant’s RDM with the RDM averaged across all train participants (lower bound). Subsequently, we squared the averaged correlation values to obtain the R^2^ values for the upper and lower bounds. We determined the unique R^2^ of each layer based on the increase in R^2^ value when that layer was included in the model.

**Figure 2.**
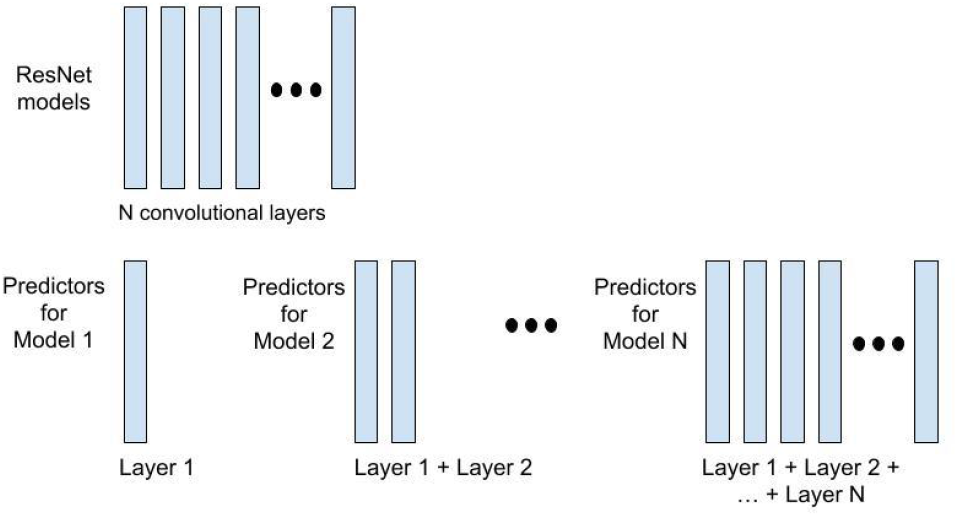
Regression models with varying number of layers as predictors. To observe the contributions of additional layers in predicting neural data, we fitted regression models with an incremental number of layers included as predictors.

### Statistical analysis: behavioral data

For all ResNets (*n*=50) and human subjects (*n*=59), categorization accuracy was computed as the average proportion of correct trials within each condition. Differences between conditions for ResNets were tested with a two-factor ANOVA (network type and background complexity), followed by *t*-tests between the condition pairs. The *p*-values obtained from the ANOVA were corrected for multiple comparisons with an FDR (α=0.01). Significance was determined based on a *p*-value which was smaller than 0.01. Behavioral analysis was performed and visualized in Python using the following packages: NumPy, SciPy, Statsmodels, Pandas, Seaborn (Harris et al., 2020; McKinney & Others, 2010; Seabold & Perktold, 2010; Virtanen et al., 2020; Waskom, 2021).

### Statistical analysis: ResNets’ explained variance on brain activity

We used a Mann-Whitney U test to test for pairwise differences in *R*^2^ between ResNets. The *p*-values obtained from the Mann-Whitney U test were corrected for multiple comparisons with an FDR (α=0.01). Similarly, the Mann-Whitney U test was used to test for differences in *R*^2^ between unmasked and masked trials for each ResNet model. Significance was determined based on a *p*-value which was smaller than 0.01.

## Results

We investigated the ability of DCNNs to capture recurrent processing in the human brain within an object categorization task. Human subjects performed the task under both visually unmasked and masked conditions. ResNets performed the recognition task with identical stimuli. We compared both the object categorization performance of human subjects with the categorization performance of ResNets, and also brain activity from human subjects with unit activations from ResNets.

### Visual masking changes human object recognition performance from one alike a deeper network into a shallower one

Human performance under visually unmasked conditions was close to performance ceiling (i.e. 100% accuracy) regardless of object background complexity (see Figure 3A). However, under visually masked conditions, human performance deteriorated with increasing background complexity. Results from repeated measures ANOVA revealed that both factors of background complexity and masking interacted – specifically, masking impaired performance to a greater degree for more complex backgrounds (Seijdel, Loke et al., 2020). For ResNets (see Figure 3B), the deepest network (i.e. ResNet-34) performed close to ceiling for all background complexity conditions; whereas shallower networks such as ResNet-10 suffered in performance as background complexity increased. The two most shallow networks - ResNet-4 and 6 - performed poorly regardless of background complexity. A two factor ANOVA was performed with network type (i.e. number of layers) and background complexity as independent factors; its results showed significant main effects of network type, *F*(4,180) = 3867.61, *p* < .001, significant main effects of background complexity conditions, *F*(3,180) = 15.93, *p* < .001, and also significant interaction effects between network type and background complexity conditions, *F*(12,180) = 7.60, *p* < .001. To examine if differences of each network’s performance across conditions were significant, pairwise comparisons *t*-tests were performed. For ResNet-4, significant differences were reported between segmented and medium complexity conditions, *t*(9) = 5.01, *p*(fdr-corrected) = .002; low and medium complexity conditions, *t*(9) = 6.00, *p*(fdr-corrected) = .001; between medium and high complexity conditions, *t*(9) = -4.19, *p*(fdr-corrected) = .005; but no significance were reported between segmented and low complexity conditions, *t*(9) = -.49, *p*(fdr-corrected) = .64; nor between segmented and high complexity conditions, *t*(9) = 1.17, *p*(fdr-corrected) = .33; nor between low and high complexity conditions, *t*(9) = 2.18, *p*(fdr-corrected) = .09.

**Figure 3.**
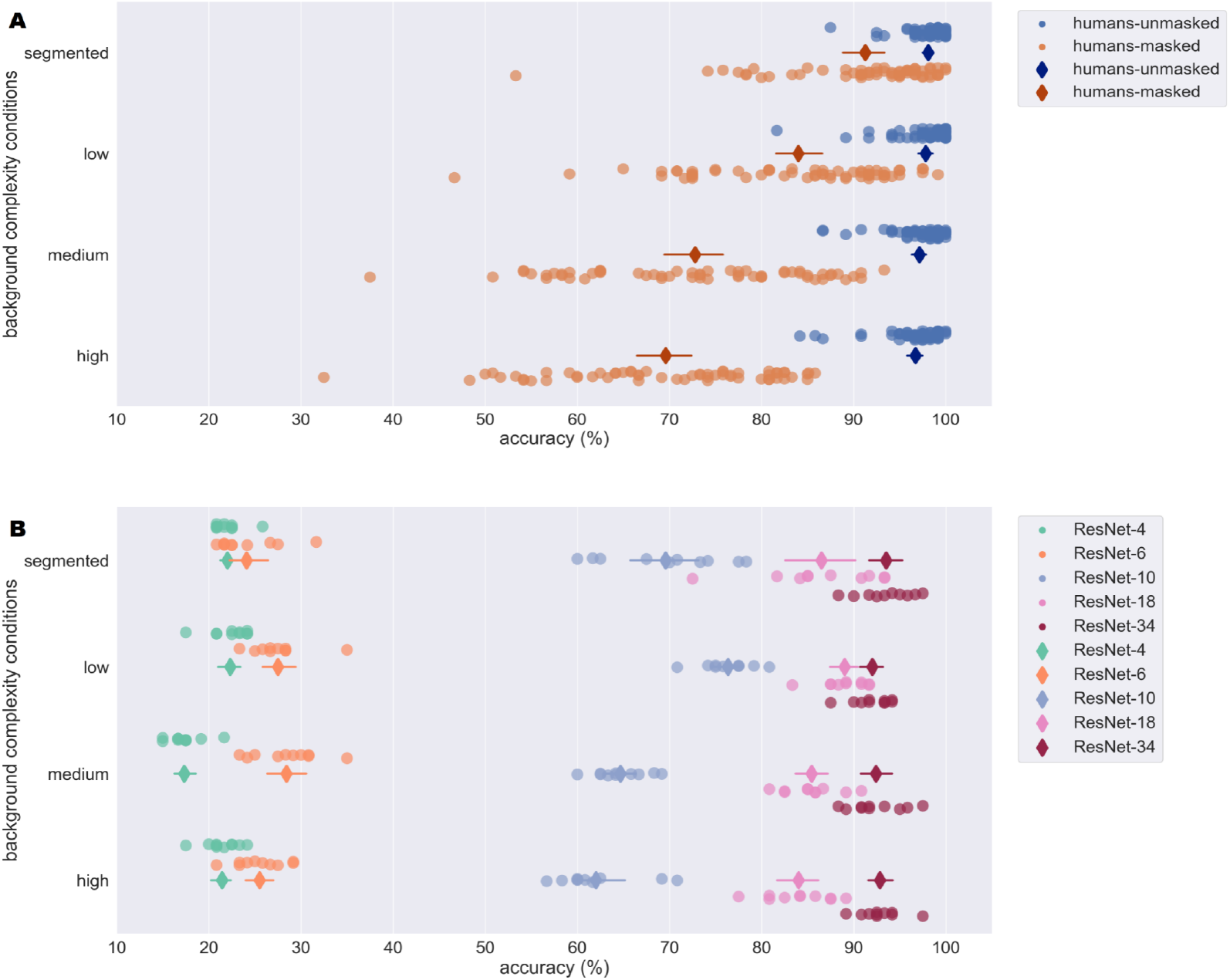
Human and ResNets’ performance on the object categorization task. Both humans and ResNets performed an object categorization task (chance performance: 20%). The different rows indicate performance in different background complexity conditions. The diamond shaped marker and line indicate average score and standard error. **A)** Human performance was consistent across background complexity conditions within unmasked trials but differed across masked trials. **B)** The deepest network (i.e. ResNet-34) mimics human performance in unmasked trials - its performance did not vary with background complexity. However, a shallower network such as ResNet-10 mimics human performance in masked trials - its performance deteriorated with medium and high complexity conditions.

For ResNet-6, significant differences were reported between segmented and low complexity conditions, *t*(9) = -4.83, *p*(fdr-corrected) = .005; but no significant differences were reported between segmented and medium complexity conditions, *t*(9) = -3.16, *p*(fdr-corrected) = .03; between segmented and high complexity conditions, *t*(9) = -0.88, *p*(fdr-corrected) = .46; between low and medium complexity conditions, *t*(9) = -.76, *p*(fdr-corrected) = .46; between low and high complexity conditions, *t*(9) = 1.30, *p*(fdr-corrected) = .34; nor between medium and high complexity conditions, *t*(9) = 2.41, *p*(fdr-corrected) = .08.

For ResNet-10, there were significant differences between low and medium complexity conditions, *t*(9) = 9.39, *p*(fdr-corrected) < .001, and between low and high complexity conditions, *t*(9) = 7.61, *p*(fdr-corrected) < .001. But no other significant differences were found - between segmented and low complexity conditions, *t*(9) = -3.08, *p*(fdr-corrected) = .02; between segmented and medium complexity conditions, *t*(9) = 2.04, *p*(fdr-corrected) = .07; between segmented and high complexity conditions, *t*(9) = 3.29, *p*(fdr-corrected) = .02; nor between medium and high complexity conditions, *t*(9) = 2.48, *p*(fdr-corrected) = .04.

For ResNet-18, no significant differences were found between all complexity conditions - between segmented and low complexity conditions, *t*(9) = -1.06, *p*(fdr-corrected) = .38; between segmented and medium complexity conditions, *t*(9) = .61, *p*(fdr-corrected) = .56; between segmented and high complexity conditions, *t*(9) = 1.27, *p*(fdr-corrected) = .38; between low and medium complexity conditions, *t*(9) = 2.83, *p*(fdr-corrected) = .06; between low and high complexity conditions, *t*(9) = 3.71, *p*(fdr-corrected) = .03; and between medium and high complexity conditions, *t*(9) = 1.22, *p*(fdr-corrected) = .38.

For ResNet-34, no significance were reported between all complexity conditions - between segmented and low complexity conditions, *t*(9) = 1.66, *p*(fdr-corrected) = .44; between segmented and medium complexity conditions, *t*(9) = 1.21, *p*(fdr-corrected) = .44; between segmented and high complexity conditions, *t*(9) = 1.12, *p*(fdr-corrected) = .44; between low and medium complexity conditions, *t*(9) = -.55, *p*(fdr-corrected) = .61; between low and high complexity conditions, *t*(9) = -1.13, *p*(fdr-corrected) = .44; and between medium and high complexity conditions, *t*(9) = -.52, *p*(fdr-corrected) = .61.

In general, each ResNet model performed consistently across different complexity conditions with the exception of ResNet-10 which performed poorer when background complexity increased from low to medium complexity.

In other words, human subjects performed optimally under visually unmasked conditions, much like a deep network (i.e. ResNet-34). But, when stimuli were masked, human subjects performed more like a shallower network (i.e. ResNet-10). Here, we can conclude that visual masking changes human object recognition performance from one alike deep networks to one alike shallower networks.

### Deeper ResNets (ResNets-10, 18 and 34) explained more variance in brain activity compared to shallower ResNets (ResNets-4 and 6)

All analyses of brain activity and ResNets activity is performed using Representational Similarity Analysis (RSA; see Materials and methods section). With RSA, we obtained stimuli pairwise dissimilarity matrix (known as representational dissimilarity matrix; RDM) as it reveals the dissimilarity distance between stimuli condition pairs. RDMs were generated based on EEG activity and network layer activations.

We tested the ability of ResNets to capture brain activity by fitting regression models for each network. In these regression models, we used ResNets RDMs as predictors for EEG RDMs. These models were fitted separately for unmasked and masked trials as we are interested to observe the difference in explained variance (*R*^2^) when recurrent processing is undisrupted (in unmasked trials) and disrupted (in masked trials). The plotted *R*^2^ for each ResNet is averaged across 10 initializations of the network. (See Figure 4)

**Figure 4.**
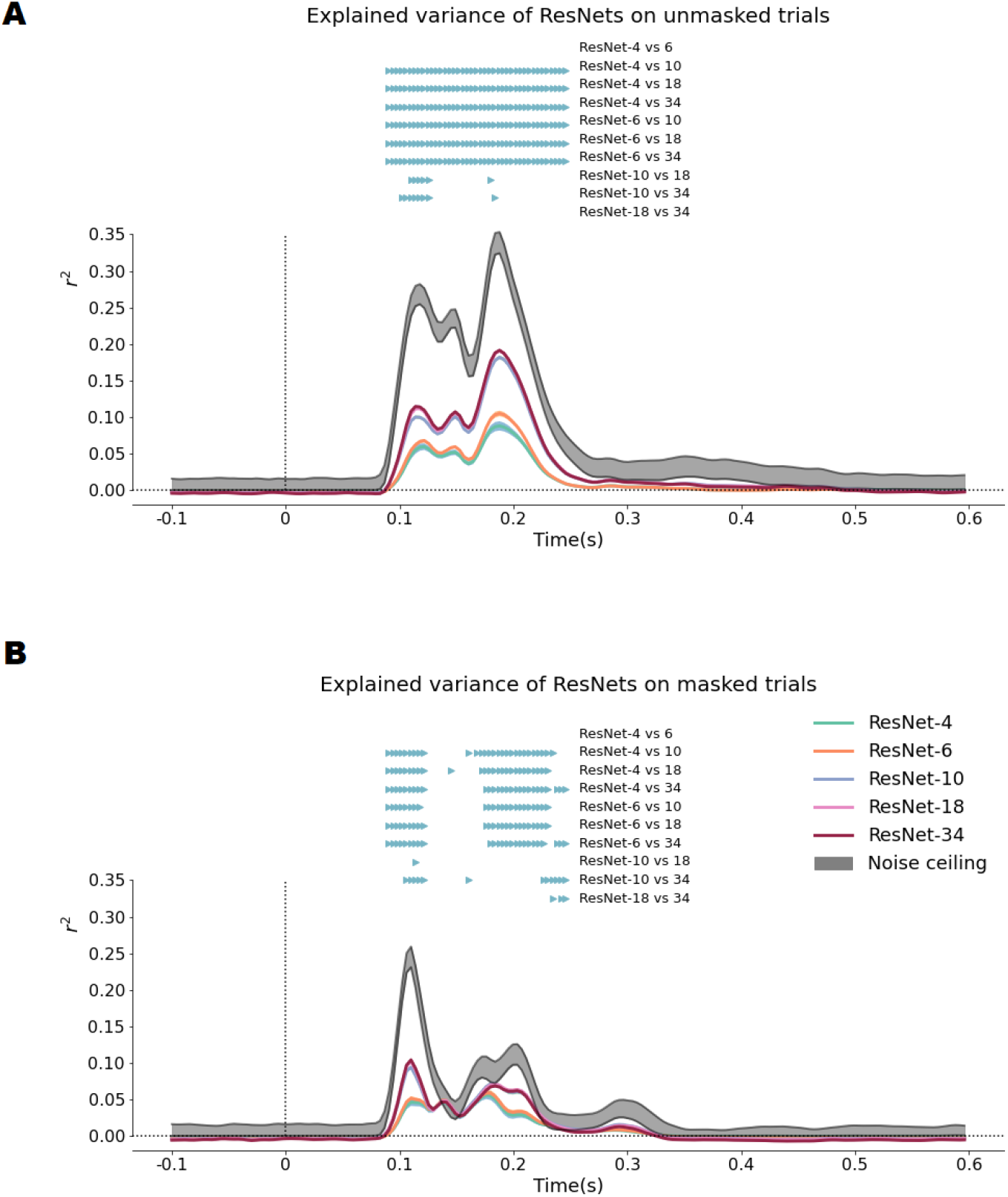
Regression models using ResNets’ layers as predictors for unmasked (A) and masked (B) trials. Pairwise comparisons between the ResNets revealed that deeper networks have higher *R*^2^ than shallower networks. **A)** Within unmasked trials, results showed that the *R*^2^ of ResNet-10, 18 and 34 is higher compared to the *R*^2^ of ResNet-4 and 6 for all time points 90–250ms. Additionally, the *R*^2^ of ResNet-18 and 34 is also higher compared to the *R*^2^ of ResNet-10. **B)** Within masked trials, results showed that the *R*^2^ of ResNet-10, 18 and 34 is higher than the *R*^2^ of ResNet-4 and 6 at two windows: 90–122ms(± 4ms) and 174-237ms(± 12ms). Additionally, the *R*^2^ of ResNet-34 is also higher compared to the *R*^2^ of ResNet-10 and ResNet-18. Surprisingly at ∼122–174ms, all networks’ *R*^2^ did not differ significantly from each other.

In order to assess if ResNet models performed differently in predicting brain activity, pairwise comparisons between ResNets were performed on samples at ∼90–250ms. Within unmasked trials, pairwise comparisons revealed no significant differences between the *R*^2^ of ResNet-4 and ResNet-6. However, significant differences were found between the *R*^2^ of ResNet-4 and 6 with the *R*^2^ of ResNet-10, 18 and 34 for all time samples within ∼90–250ms (ɑ = 0.01, FDR corrected for 420 pairwise comparisons – 10 model pairs x 42 time samples). Pairwise comparisons between the *R*^2^ of ResNet-10 and ResNet-18 have shown significant differences at ∼110–126ms, and ∼180ms (ɑ = 0.01, FDR corrected). Pairwise comparisons between the *R*^2^ of ResNet-10 and ResNet-34 have shown significant differences at ∼102–126ms, and ∼184ms (ɑ = 0.01, FDR corrected). Pairwise comparisons between the *R*^2^ of ResNet-18 and ResNet-34 revealed non-significant differences.

In summary, the *R*^2^ of ResNet-4 and ResNet-6 did not significantly differ, and both are significantly lower than the *R*^2^ of ResNet-10, 18 and 34. The *R*^2^ of ResNet-10 significantly differed from the *R*^2^ of ResNet18 at ∼110–126ms and ∼180ms. The *R*^2^ of ResNet-10 also significantly differed from the *R*^2^ of ResNet-34 at ∼102–126ms, and also ∼184ms. The *R*^2^ of ResNet-18 and ResNet-34 did not significantly differ from each other. We observed that deeper networks have higher *R*^2^ compared to shallower networks, indicating an increased capture of the variance in EEG data; this is especially apparent in the differences between ResNet-10, 18 and 34 and ResNet-4 and 6.

Subsequently, we fitted the models to masked trials. We similarly performed pairwise comparisons between the *R*^2^ of ResNet models on samples at ∼90–250ms. Pairwise comparisons between the *R*^2^ of ResNet-4 and ResNet-6 revealed no significant differences. Pairwise comparisons between the *R*^2^ of ResNet-4 and 6 with ResNet-10, 18 and 34 significantly differed in two windows: ∼90–122ms (± 4ms) and ∼174–237ms (± 12ms), (ɑ = 0.01, FDR corrected). Pairwise comparisons between the *R*^2^ of ResNet-10 and ResNet-18 revealed only one significant time sample at ∼114ms (ɑ = 0.01, FDR corrected). Pairwise comparisons between *R*^2^ of ResNet-10 and ResNet-34 showed significant differences at ∼106–122ms, ∼161ms, and also ∼227–246ms (ɑ = 0.01, FDR corrected). Pairwise comparisons between *R*^2^ of ResNet-18 and ResNet-34 showed significant differences at ∼234ms and ∼242–246ms (ɑ = 0.01, FDR corrected). However, *R*^2^ of all ResNets did not significantly differ from each other at ∼122–174ms.

In summary, the *R*^2^ of ResNet-4 and ResNet-6 for masked trials do not show significant differences. However, the *R*^2^ of ResNet-10, 18 and 34 is higher than the *R*^2^ of ResNet-4 and 6 at ∼90ms–122ms (± 4ms) and ∼174–237ms (± 12ms). Only one significant time point was found between between *R*^2^ of ResNet-10 and ResNet-18 at ∼114ms. Comparisons between *R*^2^ of ResNet-10 and ResNet-34 showed significant differences at ∼106–122ms, ∼161ms, and also ∼227–246ms. Comparisons between *R*^2^ of ResNet-18 and ResNet-34 showed significant differences at ∼242–246ms. Similar to unmasked trials, we also observed that deeper networks (ResNet-10, 18 and 34) have higher *R*^2^ compared to shallower networks (ResNet-4 and 6); however, the duration of significant differences between both deeper and shallower networks have shrunk in masked trials compared to unmasked trials. Additionally, within masked trials, the *R*^2^ of all ResNets did not significantly differ from each other at∼122–174ms.

To compare between the explained variance in unmasked and masked trials, we performed pairwise comparisons between the models’ *R*^2^ for unmasked trials and *R*^2^ for masked trials on time samples ∼90–250ms. For ResNet-4, significant differences were found – ∼122–129ms, ∼145–157ms, ∼176–231ms, and ∼246ms (ɑ = 0.01, FDR corrected). For ResNet-6, significant differences were found at ∼118–133ms, ∼145–157ms, and ∼172–231ms (ɑ = 0.01, FDR corrected). For ResNet-10, significant differences were found at ∼102–106ms, and ∼114–246ms (ɑ = 0.01, FDR corrected). For ResNet-18, significant differences were found at ∼98–106ms, and ∼114–246ms (ɑ = 0.01, FDR corrected). For ResNet-34, significant differences were found at ∼102–106ms, and ∼114–246ms (ɑ = 0.01, FDR corrected).

In short, significant differences in *R*^2^ between masking conditions were found as early as ∼98ms for ResNet-18, ∼102ms for ResNet-10 and 34, ∼118ms for ResNet-6 and ∼122ms for ResNet-4. There were no effects of masking nor network depth before ∼98ms, indicating EEG activity reflected purely feedforward processes.

Additionally, we computed the difference between the *R*^2^ for unmasked trials and masked trials (see Figure 5B) and observed that the difference between masking conditions gradually increased to a peak at ∼200ms. The magnitude of differences between masking conditions are also larger for deeper networks (ResNet-10, 18, and 34) than shallower networks (ResNet-4 and 6), indicating that deeper networks are better at capturing recurrent processing signals at later time-points.

**Figure 5.**
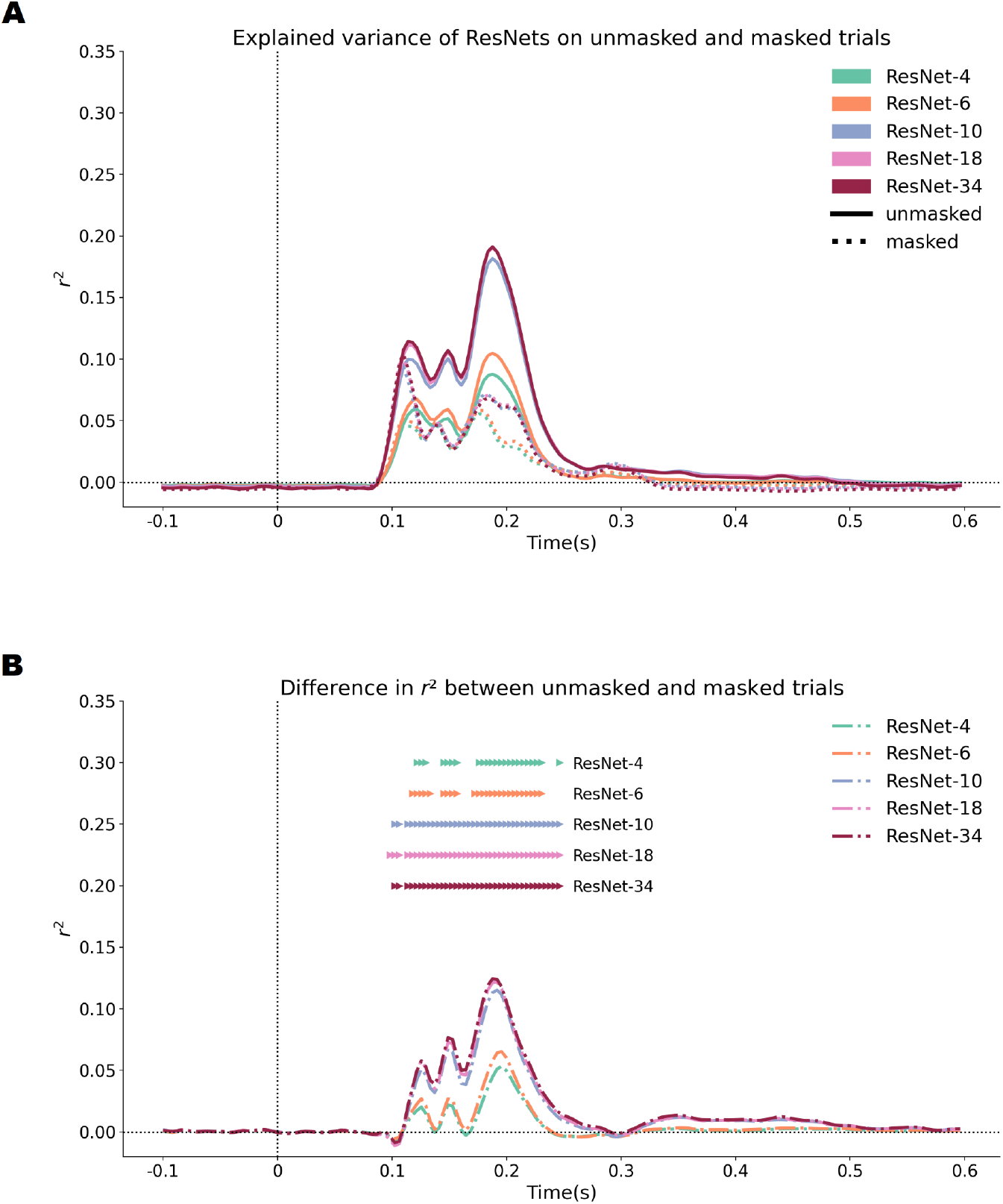
R^2^of models for unmasked and masked data and differences in models performance between masking conditions. **A)** The *R*^2^ for both unmasked and masked trials are plotted together. **B)** The difference in *R*^2^ between masking conditions is plotted. Colored markers above reflect significant differences between masking conditions. We similarly see that deeper networks (ResNet-10, 18 and 34) have longer duration of significant differences compared to shallower networks (ResNet-4 and 6), moreover, the magnitude of differences are larger for deeper networks.

### Early DCNN layers are sufficient to explain brain activity

To understand the contribution of ResNets’ depth in capturing brain activity, we built a series of regression models differentiated by the number of convolutional layers used as predictors (see Figure 2). As more layers are included as predictors, the model increases in *R* ^2^ (see Figure 6). However, at a certain number of layers, including more layers no longer increases the *R*^2^ of the model. ResNet-4 reached maximal ^2^ in layer 3. ResNet-6 and ResNet-10 reached maximal *R*^2^ in layer 5. ResNet-18 reached maximal *R*^2^ in layer 7. ResNet-34 reached maximal *R*^2^ in layer 9. Deeper models like ResNet-10, 18 and 34 reached maximal *R*^2^ in its early layers (i.e. first half layers of the network). Layers beyond these layers of maximal *R*^2^ do not improve the model performance (i.e. amount of *R*^2^).

**Figure 6.**
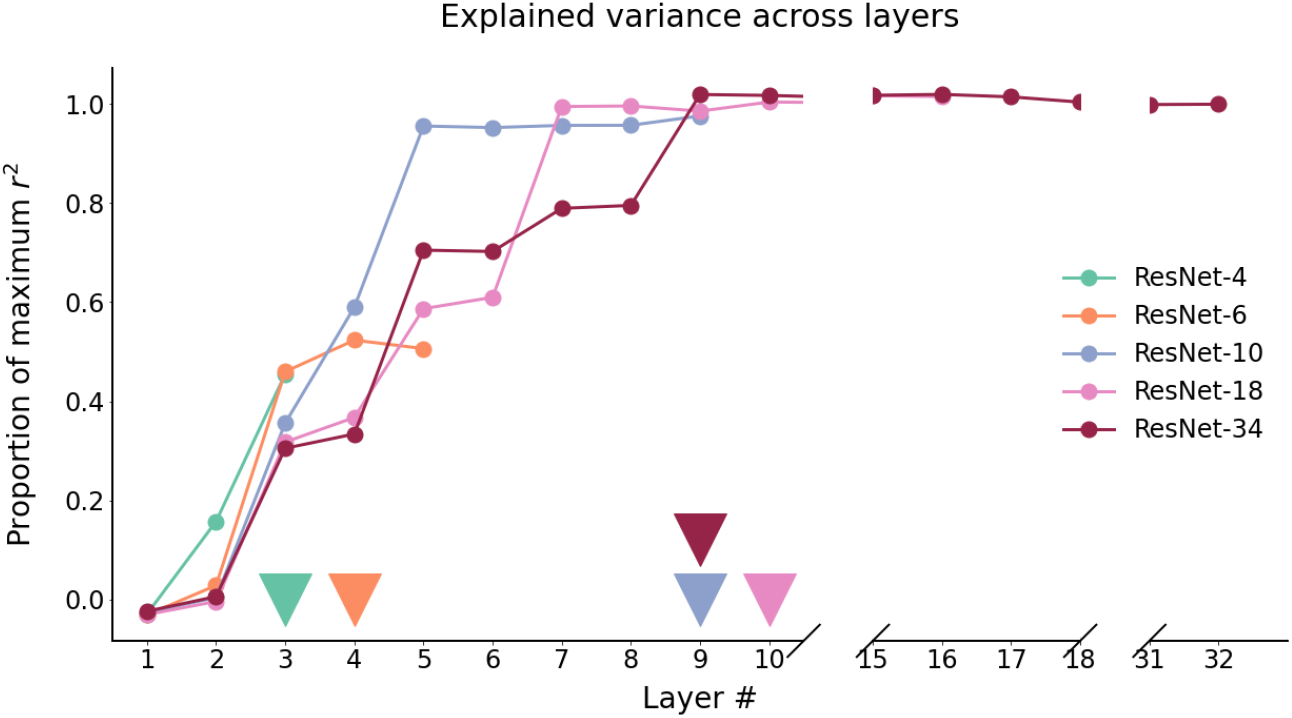
Models’ R^2^ plotted as a proportion of the maximum R^2^. We fitted regression models with an increasing number of convolutional layers as predictors. With each additional layer, models increases its *R*^2.^ However, the increase in *R*_2_ stopped at a certain number of layers. The colored triangles indicate the layer where *R*^2^ stopped increasing for the model. For ResNet-10, 18 and 34, the maximal *R*^2^ was reached in its early layers (i.e. first half layers of the network).

To further assess the networks’ ability to predict brain activity, we computed the unique *R*^2^ from each network layer to observe each layer’s contribution to the model performance. The unique *R* ^2^ of a layer is computed as the increase in *R*^2^ of the model with the additional layer as a predictor (See equation below).

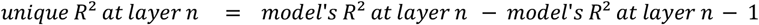

We observed that the early layers of deeper networks contributed disproportionately to the models’ *R*^2^(see Figure 7). Layers 3, 4 and 5 contributed most to deeper networks’ (ResNet-10, 18 and 34) *R*^2^. Paradoxically, though deeper networks have significantly higher *R* ^2^than shallower networks, deeper networks’ performance can be attributed to activity in its early layers; deeper layers (i.e. second half layers of the network) do not further contribute to the models’ *R*^2^.

**Figure 7.**
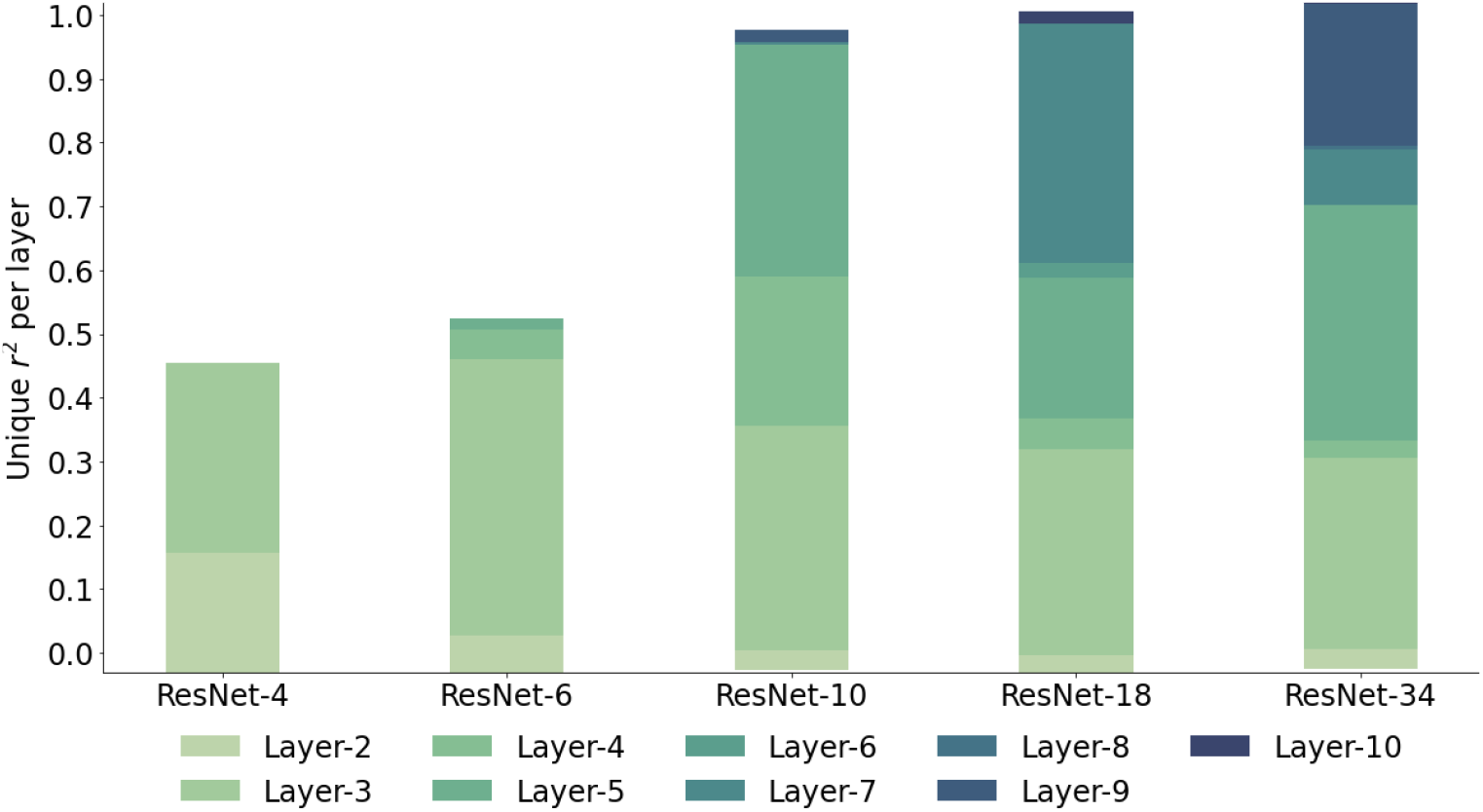
Unique R^2^ of each ResNet layer. We computed the unique variance for each layer in the regression model. For shallower networks (ResNet-4 and 6), layers 2 and 3 contributed most to the model. For deeper networks (ResNet-10, 18 and 34), layers 3, 4 and 5 similarly contributed at least half of the models’ *R* ^2^. Note: Layer-1 is not plotted as it has negative *R*^2^ values which happen when the model is arbitrarily worse than chance. This is also the reason why the plot begins below zero.

## Discussion

### Summary

In this study, we investigated the ability of DCNNs to capture recurrent processing in the human brain. Specifically, we tested ResNets as they approximate an additive form of recurrent processing consisting of repeating excitatory activations on static inputs. We used ResNets of varying depths as proxies for varying amounts of recurrent processing. We expected deeper networks (i.e. networks with more recurrent processing) to explain more variance in brain activity (i.e. have higher *R*^2^) than shallower neworks in unmasked trials when recurrent processing was not disrupted, but for all networks to have similar *R*^2^ in masked trials when recurrent processing was disrupted. Our expectations were partially met as deeper networks (ResNet-10, 18 and 34) indeed have higher *R*^2^ than shallower networks (ResNet-4 and 6), but for both unmasked and masked trials. We also find that all ResNet models (ResNet-4, 6, 10, 18, 34) have higher *R*^2^ under unmasked trials than masked trials, with differences in *R*^2^ starting as early as ∼98ms. These differences in *R*^2^ gradually increased to a peak at ∼200ms, with deeper networks (ResNet-10, 18, 34) showing larger magnitudes of differences compared to shallower networks (ResNet-4 and 6). By building regression models with increasing numbers of layers as predictors, we see that only early layers (i.e. first half layers of the model) contributed to the *R*^2^ of the model.

### Deeper networks capture behavioral performance but not recurrent processes in early visual cortex

We found that humans performed similarly as a deep network (i.e. ResNet-34) under visually unmasked conditions; but visual masking deteriorated human performance to become more like a shallower network (i.e. ResNet-10). Based on categorization as performance, we had expected a deep model like ResNet-34 to have higher *R*^2^compared to a shallower model like ResNet-10. This expectation stems from previous studies - where DCNNs that perform better in categorization accuracy also better predict brain data (Yamins et al., 2014). However, although ResNet-34 significantly outperforms ResNet-10 at categorizing objects, the *R*^2^ ResNets-34 only significantly differed from the *R*^2^ of ResNet-10 for a short time window at ∼102–126ms and ∼184ms. Nonetheless, our finding agrees that model depth improves the model’s object categorization performance - similar to the findings of the original ResNet creators (He et al., 2016a). Though, the mechanisms in ResNets’ deeper layers do not seem to match the underlying mechanisms in human’s early visual cortex, at least as measured with EEG. This discrepancy between behavioral outcomes and brain activity prediction can also be observed on the Brain-Score platform, where we see that the unmodified version of ResNets tend to score well on explaining behavior but score less well on explaining activity in V1 (Schrimpf et al., 2018).

### Simple recurrence captures recurrent processing but still insufficient

ResNets are made up of residual blocks. Each residual block has multiple convolutional layers with the same filter size and same number of filters. Thus, the repetition of identical convolutional operations could be perceived as an additive form of recurrence - recurrence by repeating excitatory activations in response to a static stimulus, similar to recurrence in current RNNs and DCNNs like CorNet. We observed that this simple form of recurrence indeed contributed to significant differences between ResNet-4 and 6 with ResNet-10, 18 and 34. However, this form of recurrence only works to a certain extent. With deeper networks, our results revealed that the *R*^2^ of ResNets-10, 18 and 34 only significantly differed on a few time points (between ResNet-10 and 34) or none at all (between ResNet-18 and 34), while there was still a large gap between ResNet-34’s *R*^2^ with the noise ceiling. Thus, it appears that though additive recurrence captures recurrent processing partially, by itself it is insufficient to fully capture recurrent processing in humans. This finding is supported by studies in animals demonstrating the importance of inhibition in feedback processes (Klink et al., 2017), and also recent studies modeling recurrent signals, where researchers have shown that both lateral (i.e. repeating excitatory activations) and feedback (i.e. top down) connections are necessary to improve the model’s performance beyond feedforward processes (Kietzmann, Spoerer, et al., 2019; Spoerer et al., 2017).

### Recurrent signals set in as early as ∼98ms

When we observed the difference in *R*^2^ between masking conditions, we see significant differences as early as ∼98ms for ResNet-18, ∼102ms for ResNet-10 and 34, ∼118ms for ResNet-6 and ∼122ms for ResNet-4 – with deeper networks (ResNet-10, 18 and 34) showing earlier significant differences compared to shallower networks (ResNet-4 and 6). These early differences indicate that recurrent signals set in as early as ∼98ms. Before ∼98ms, there were no effects of masking nor effects of network depth, indicating that EEG activity before ∼98ms are feedforward processes. The differences in *R*^2^ between masking conditions gradually increased until a peak difference at ∼200ms, indicating that recurrent signals build up across time. As such, early EEG activity is a mixture of feedforward and recurrent processes, whereas late EEG activity is mainly dominated by recurrent processes. Consequently, our results suggest that models including interactions between feedforward and feedback streams across time steps could better capture recurrent processes in humans.

## Conclusion

In this paper, we tested the ability of DCNNs to capture recurrent processes in human brains. Specifically, we tested an additive form of recurrence in ResNets to predict recurrent signals in human visual systems. We found that deeper ResNets (ResNets-10, 18 and 34) explained more variance in brain activity than shallower ResNets (ResNets-4 and 6). Furthermore, all ResNets explained more variance in brain activity during unmasked trials than masked trials. Differences in explained variance between masking conditions set in as early as ∼98ms and gradually increased to a peak at ∼200ms, indicating that early brain activity consists of both feedforward and recurrent processes but gradually becomes dominated by recurrent processes. Accordingly, deeper networks showed larger differences in explained variance between masking conditions than shallower networks, providing further evidence that deeper networks capture larger proportions of recurrent processing signals. However, given the substantial distance between the models’ explained variance and data’s noise ceiling, we posit that other types of recurrent processes (inhibition, multiplicative), that are not present in current regular deep neural networks (alexnet, cornet, resnet) are of paramount importance towards better visual models.

## Acknowledgements

This work is supported by an Interdisciplinary Doctorate Agreement from the University of Amsterdam to H. Steven Scholte and Natalie Cappaert and an Advanced Investigator Grant from the European Research Council (ERC) to Edward de Haan (#339374).

## Notes

**Conflict of interest:** The authors declare no competing financial interests.

### Competing Interest Statement

The authors have declared no competing interest.

